# Environmental dependence of colony morphologies in *Labyrinthula* species

**DOI:** 10.1101/2025.04.01.646626

**Authors:** Joseph M. Knight, Patricia Gonzalez, David J. Fairhurst, Gavin Melaugh, Wilson C. K. Poon

## Abstract

*Labyrinthula* species are protist organisms found across a variety of marine environments whose defining characteristic is the secretion of an extracellular ectoplasmic net. Under certain conditions, colonies form a spatial network of ‘tracks’ through which cells move bidirectionally. We show that this network morphology depends on the presence of a liquid overlay, with air exposed colonies exhibiting instead a dense, aggregated morphology. We demonstrate dynamic restructuring between these two morphologies upon addition or removal of the liquid overlay, and investigate growth behaviour under varying nutrient conditions. Given the inter-tidal environment of certain seagrass species colonised by *Labyrinthula*, our results may shed light on the relationship between this organism and its seagrass host, for which it is an opportunistic pathogen associated with seagrass wasting disease.

## INTRODUCTION

Coastal regions comprise 4% of the world’s land surface area, but are home to one third of its population [1]. Seagrasses, a group of over 70 species of marine flowering plants [2] that includes the oldest known plant on Earth [3], act as vital ‘ecosystem engineers’ [4] to create, modify and maintain coastal habitats, supporting biodiversity at multiple trophic levels [5]. By capturing carbon dioxide for their own photosynthesis, collecting suspended organic matter, and stabilising sediment, coastal seagrass meadows are also some of the most efficient carbon sinks on earth [6]. While many species such as the common eelgrass (*Zostera marina*) are predominantly submerged, there are also inter-tidal species such as the dwarf eelgrass (*Z. noltii*) that are regularly exposed to air at low tide.

Today, the global seagrass population is in decline [7], due at least in part to climate change [8], with some 10 species at elevated risk of extinction [9]. Previously, a global decline occurred in the early 1930s, when an epidemic of ‘wasting disease’ virtually eliminated eelgrass from both sides of the Atlantic [10], although strangely, *Zostera noltii* was largely unaffected [11]; smaller episodes have recurred thereafter [12].

Following the epidemic of the 1930s, the disease became associated with the unicellular protist *Labyrinthla* spp. [13], cells of which have been repeatedly isolated from characteristic dark, necrotic lesions on *Z. marina* leaves [10, 14, 15]. It is now known that pathogenic isolates of *Labyrinthula* are endemically present in eelgass but need not cause epidemic-scale depopulation [16], and non-virulent infection can lead to faster-growing (but fewer) leaves [17]. Moreover, *Labyrinthula* is a highly-connected node in the seagrass microbiome [18], reflecting, e.g., the fact that it feeds on diatoms [19], which are ubiquitous in *Zostera* beds [20]. So, *Labyrinthula* is perhaps an opportunistic pathogen of seagrass [21], although ‘pathobiont’ [22] may be more appropriate for a relationship that appears to be ‘at the edge between physiology and pathology’ [23]. Irrespective of nomenclature, the interaction is complex [24].

This complex picture begs a simple question: what triggered the endemically-present *Labyrinthula* to cause a catastrophic epidemic in *Z. marina* in the 1930s and subsequent, less severe, episodes [10]? The question is urgent in the context of global seagrass population decline [7, 9] and coastal environment degradation [8]. Potential triggers such as temperature [25, 26] and salinity [27] have been studied individually in the laboratory, but elucidating their interaction *in natura* requires advances on many fronts, including a better understanding of *Labyrinthula* biology.

*Labyrinthula*, first described in 1867 by Cienkowski [29], belongs to a clade of ‘fungal-like organisms’, the Stramenopiles [30] (Latin *stramen* = straw, *pilus* = hair) Fig. 1, many of which are unicellular flagellates. They typically bear hair-like extensions (mastigonemes) on an anterior flagellum at some stage of their life cycle, which in *Labyrinthula* corresponds to the asexual gamete (or zoospore) phase [31]. In the phylum Bigyra, each flagellum is joined to the cell body by a double helical structure (Greek *gyros* = turn) [32]. The class Labyrinthuomycota is distinguished by the existence of a membrane-bound extracellular ectoplasmic network of tracks for colonising their substrates. The class name alludes to fungal mycelia, while its popular name, the marine slime moulds, alludes to the plasmodium of acellular organisms such as *Physarum polycephalum* [33], although these organisms are neither fungi nor slime moulds.

**FIG. 1.**
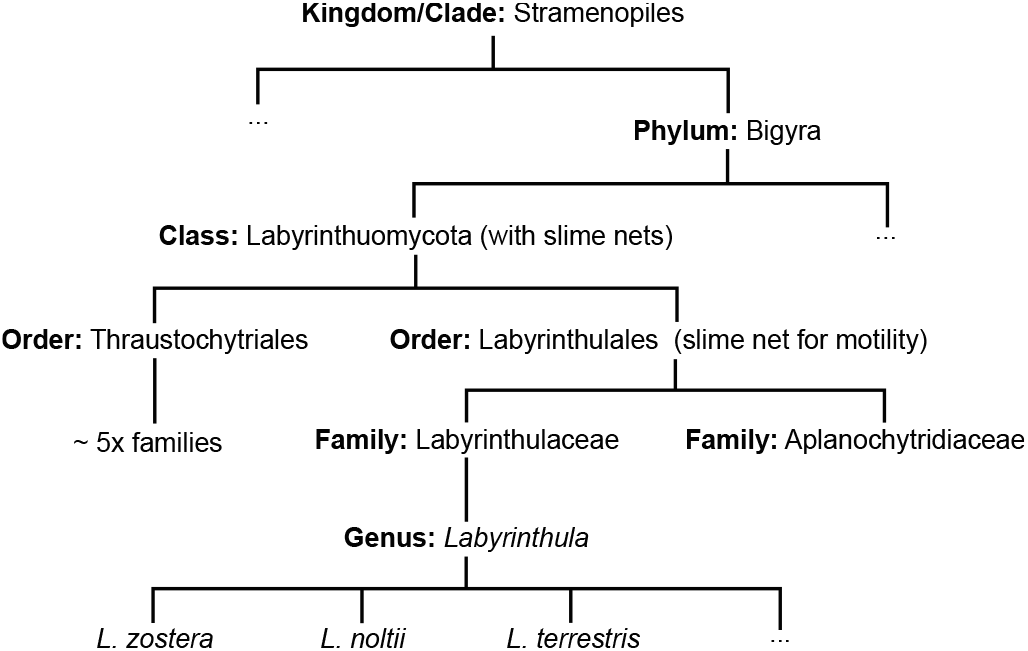
The taxonomy of *Labyrinthula*.

The Labyrinthuomycota contains two orders, the Labyrinthulids and the Thraustochydrids [13, 30]. The latter, presumably so named [34] because spores are released from bursting spherical sporangia (Greek *thraustos* = fragile, *chytridion* = little pot), use their slime nets for osmotrophic feeding, secreting extracellular digestive enzymes and absorbing and transporting nutrients [35, 36]. Genomics suggests that Labyrinthulid slime nets are also involved in feeding [37]. Additionally however, cells move bidirectionally along the trackways [38] [39], representing a gain of function from using the net for nutrition alone [40].

Figure 2(a) shows a putative scheme for the initial stages of *Labyrinthula* colony formation on a substrate from a single cell [28, 41]. A membrane organelle, the bothrosome (Greek *bothros* = hole, *soma* = body) [41], secretes a membrane ‘bubble’ that expands and envelops the cell. Membrane fusion then forms two concentric extracellular membranes that enclose an ectoplasm, which completely surrounds the cell. Cell division generates further cells inside the outer ectoplasmic membrane.

**FIG. 2.**
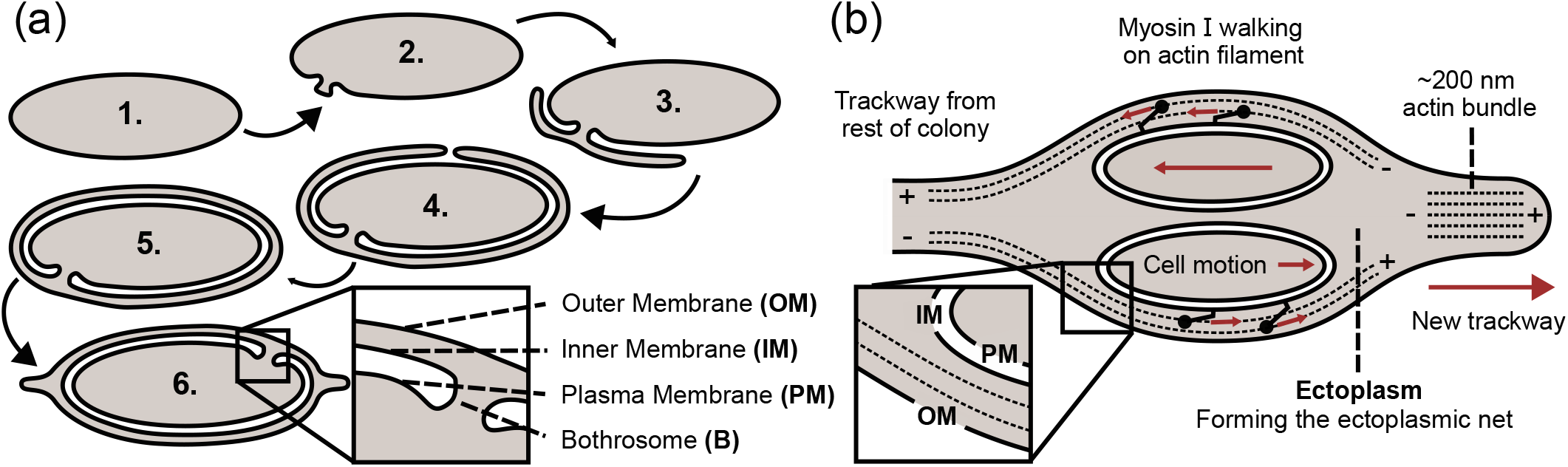
*Labyrinthula* schematics. (a) Putative early stages of *Labyrinthula* colony formation from a single cell showing the role of the bothrosome in extracellular membrane production (adapted from [28]). (b) Putative arrangement of membranes, actin filaments (polarity indicated by + and -) and myosin I motors (filled circles with ‘tails’) in *Labyrinthula*. The different components are not drawn to scale.

Actin is ubiquitously present throughout the ectoplasm, although its mode of biogenesis remains obscure [42]. At the edge of the ectoplasmic matrix, actin polymerises to form filopodia-like protrusions consisting of ≳ 200 nm-thick bundles of actin filaments (individually ∼7 nm thick) [42], Fig. 2(b).

The actin bundles are visible in optical microscopy as the ‘slime net’ already noted by Cienkowski [29]. The tracks do not rest directly on substrates, but can be ‘anchored’ by protruding lateral filaments substantially thinner than the tracks that act as ‘guy ropes’ [42].

ATP-powered single-headed myosin I motors ‘walking’ on filaments likely drive cellular motility [42, 43]. Electron and optical microscopy show that cells move through (rather than on) the actin bundles, temporarily expanding each bundle into parallel strands of individual actin filaments [42], Fig. 2(b). Bidirectional motion implies filaments of both polarities, although the nature of the coordination between multiple motors to enable such motion, e.g., through some form of strain-dependent cooperativity [44], is yet to be elucidated.

Different submerged habitats for *Labyrinthula* spp. have been reported from early on [45, 46]. Their ectoplasmic network has therefore not been specifically adapted for colonising eelgrass and for its transmission by leaf-to-leaf contact [47]. Moreover, *Labyrinthula* is able to colonise *Z. noltii* [11], which is periodically exposed to air at low tides. Indeed, even drier conditions can be tolerated by *L. terrestris*, which causes ‘rapid blight’ and death in turfgrass [48] and has been isolated from other land grass species [49]. *Labyrinthula* spp. are therefore versatile colonisers of many substrates under a range of conditions [13].

As a prelude to understanding these diverse natural habitats, we have undertaken a systematic study of how a species isolated from *Z. marina* colonises standard laboratory substrates – hard, non-porous plastic and soft, porous agar gel – under submerged and dry conditions. We first report the extent of growth and gross morphology, and place these findings in the context of *Labyrinthula* biology *in natura*. Next, we explore how an ectoplasmic network emerges from small clusters of single cells. We observe twisting and buckling of filaments at the early stages of colony development that is reminiscent of similar motion in the filopodia of human cells [50, 51], suggesting that aspects of *Labyrinthula* biology may find parallel in other organisms. Finally, we map how a mature colony restructures, finding evidence for a potential role of cells at the colony edge in expanding the ectoplasmic network.

## MATERIALS AND METHODS

We used a *Labyrinthula* strain isolated from *Z. marina* harvested from Tyninghame Bay, East Lothian, Scotland. Collected leaves were washed in ethanol, cut into 1 cm strips and placed on seawater serum agar (SSA). To prepare 1 L of SSA, 12 g agar, 1 g glucose, 0.1 g yeast extract and 0.1 g peptone (all Sigma-Aldrich) were dissolved in 1 L of InstantOcean artifical seawater (33 g InstantOcean sea salt in 1 L Milli-Q water ≡ 33 PSU salinity). Following 15 min autoclaving at 121 ^°^C, 25 mL Penicillin-Streptomycin (Gibco) and 10 mL horse serum (Gibco) were added before plates were poured. Axenic cultures of *Labyrinthula* were maintained by weekly transfer of cells onto fresh SSA plates. For liquid overlays, a seawater serum broth (SSB) was prepared as SSA but with the exclusion of agar. InstantOcean artificial seawater was also used on its own as a liquid overlay. 20 mL of the liquid was transferred to the inoculated agar plate to create the overlay. Apart from observing colonies on the macro-scale on agar plates, we also used flowcells to image the microscopic behaviour of *Labyrinthula* colony development. In the latter case, cells were harvested from axenic culture plates and inoculated into 1 mL SSB before transfer to the flowcell (Luer μ-Slide, Ibidi). In all cases, imaging was performed using an inverted phase contrast microscope with a 10×, air objective.

## RESULTS AND DISCUSSION

### *Labyrinthula* colony morphology is dependent on a liquid overlay

We find that the morphology of *Labyrinthula* colonies on agar depends dramatically on the environmental condition above the agar. Following inoculation onto Seawater Serum Agar (SSA), the addition of a seawater liquid overlay results in colonies expanding as sparse, filamentous networks, Fig. 3(a)-(b). This network morphology, characteristic of the order, displays an archetypal ectoplasmic slime net within which the spindle cells are confined to move, Fig. 3(c). Strikingly, network structures are visible even on the macro-scale, Fig. 3(a), as radial trackways emanating from the initial inoculum. Exploratory filopodia can be seen at the colony front, Fig. 3(c), driving colony expansion.

**FIG. 3.**
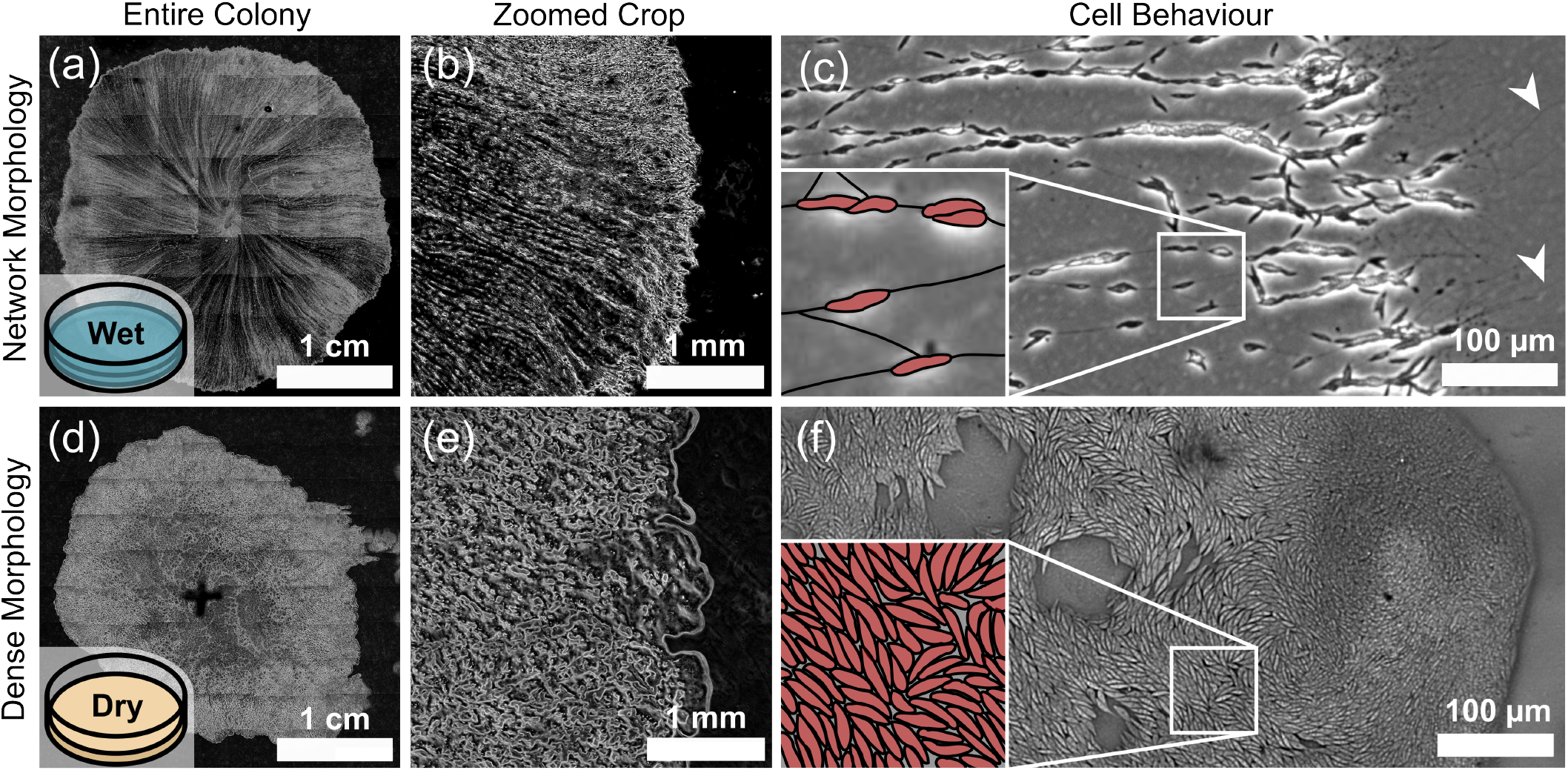
Wet and dry morphologies of *Labyrinthula*. (a)-(b) *Labyrinthula* colony inoculated on SSA with a seawater overlay showing the filamentous network morphology after twenty hours of growth. (c) Association of spindle cells (segmented and highlighted red in the inset) to the ectoplasmic net filaments visible in the colony bulk, and at the expanding front behaving as exploratory filopodia (see arrows). (d)-(e) Without a liquid overlay, the dry colony takes around five days to reach a similar spatial extent shown in (a), exhibiting the densely packed morphology. (f) Dense packing of cells at the colony front (segmented and highlighted in red in the inset). Captured in phase contrast (except (f), which was captured in brightfield) using a 10× air objective, with parts (a) and (b) being grids of 10 × 10 tiled images.

In stark contrast, without a liquid overlay, inoculated *Labyrinthula* exposed directly to air grow as a densely-packed colony with occasional voids, Fig. 3(d)-(f). In this dense state, it is difficult to resolve the ectoplasmic net typically associated with the spindle cells, and therefore how the putative membrane scheme of the network morphology (Fig. 2(b)) applies to this phase. However, we observe cell motion in this dense phase (Electronic Supplementary Video 1) that is reminiscent of the motion observed previously [52] in smaller membrane-bound ‘whirling aggregates’, so that an entire dense colony could also be similarly enclosed. At the expanding front, we do not observe the exploratory filopodia characteristic of the network morphology, with cells instead typically aligned parallel to the front. This morphology is reminiscent of that seen in rod-like *Escherichia coli* bacteria growing on agar, in which cell division drives the outwards expansion of a colony of densely-packed cells [53]. By contrast, when a liquid overlay is present, cell motility drives the outward expansion of a sparse network of trackways.

### Nutrient presence determines colony spatial extent

To explore further the environment dependence of colony morphology, we inoculate *Labyrinthula* on substrates of varying composition and with different overlays. Figure 4 reports the colony morphology and area after 20 hours of growth for different substrate-overlay pairs. With no nutrients, the initial inoculum does not expand. With nutrients in either the substrate (SSA) or liquid overlay (SSB), *Labyrinthula* is able to expand and colonise the substrate surface. When nutrients are present in the liquid overlay (SSB), little substrate dependence is observed, Fig. 4(b) (compare all ‘X + SSB’). However, each of these environments gives rise to a larger colony after 24 hours than for a seawater overlay on nutritive substrate (SW + SSA). These observations suggest faster or preferential nutrient uptake from the overlying liquid than from an underlying solid substrate. The reason for this is currently unclear. Slower diffusion of nutrients through our 1.2% agar is not the cause: glucose [54], globular proteins [55] and lipids [56] have comparable diffusivities in 1-3% agar and water. Faster non-diffusive transport, such as advective flows in the liquid overlay may explain this observed discrepancy.

**FIG. 4.**
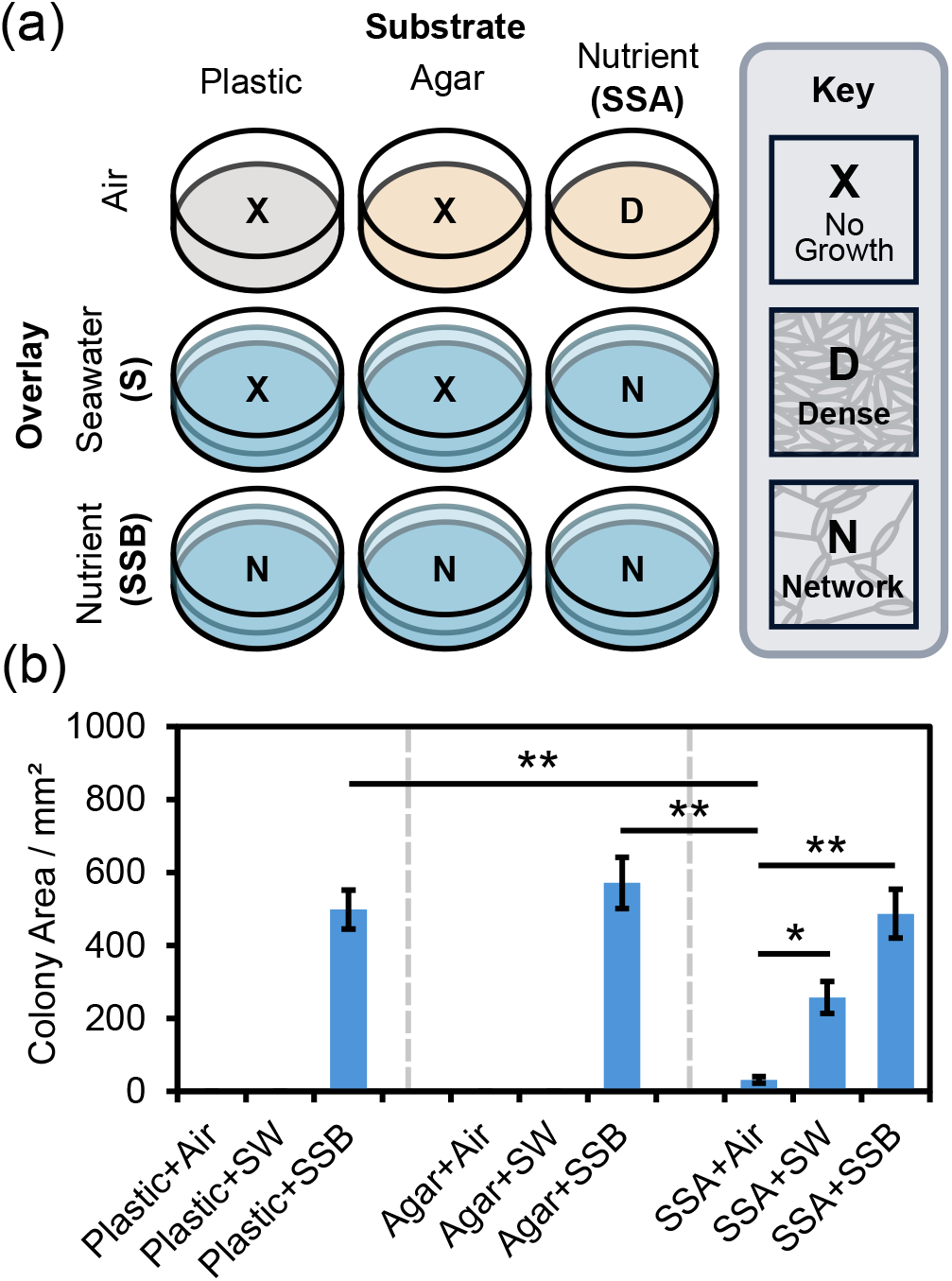
Substrate and Immersion Variation. (a) Dependence of colony morphology on substrate and immersion medium. (b) Spatial extents of colonies after 20 h of growth with data shown as mean ± SEM (n=6). Paired student’s *t*-test was used; \**p* < 0.01, \*\**p* < 0.001.

### Addition and removal of liquid overlay switches colony morphology

Given the distinct morphological states under wet (submerged) or dry (exposed to air) conditions, Fig. 3, we explored whether a transition between these states could be induced in real time by a sudden change in the environment by adding or removing seawater to dry and wet colonies respectively. Submerging under seawater a colony initially growing at the agar-air interface leads to a transition from a dense to a sparser, network morphology within minutes, Fig. 5(a). Similarly, for a submerged colony showing the network morphology, removal of the seawater overlay results in a switch to the dense, aggregated structure, Fig. 5(b). We tracked the front expansion across the transition in each case, Fig. 6, and find that the air-exposed colonies expand radially at a slower rate (∼1 µm min^−1^) than submerged, network colonies (∼ 10 µm min^−1^), consistent with Fig. 4(b).

**FIG. 5.**
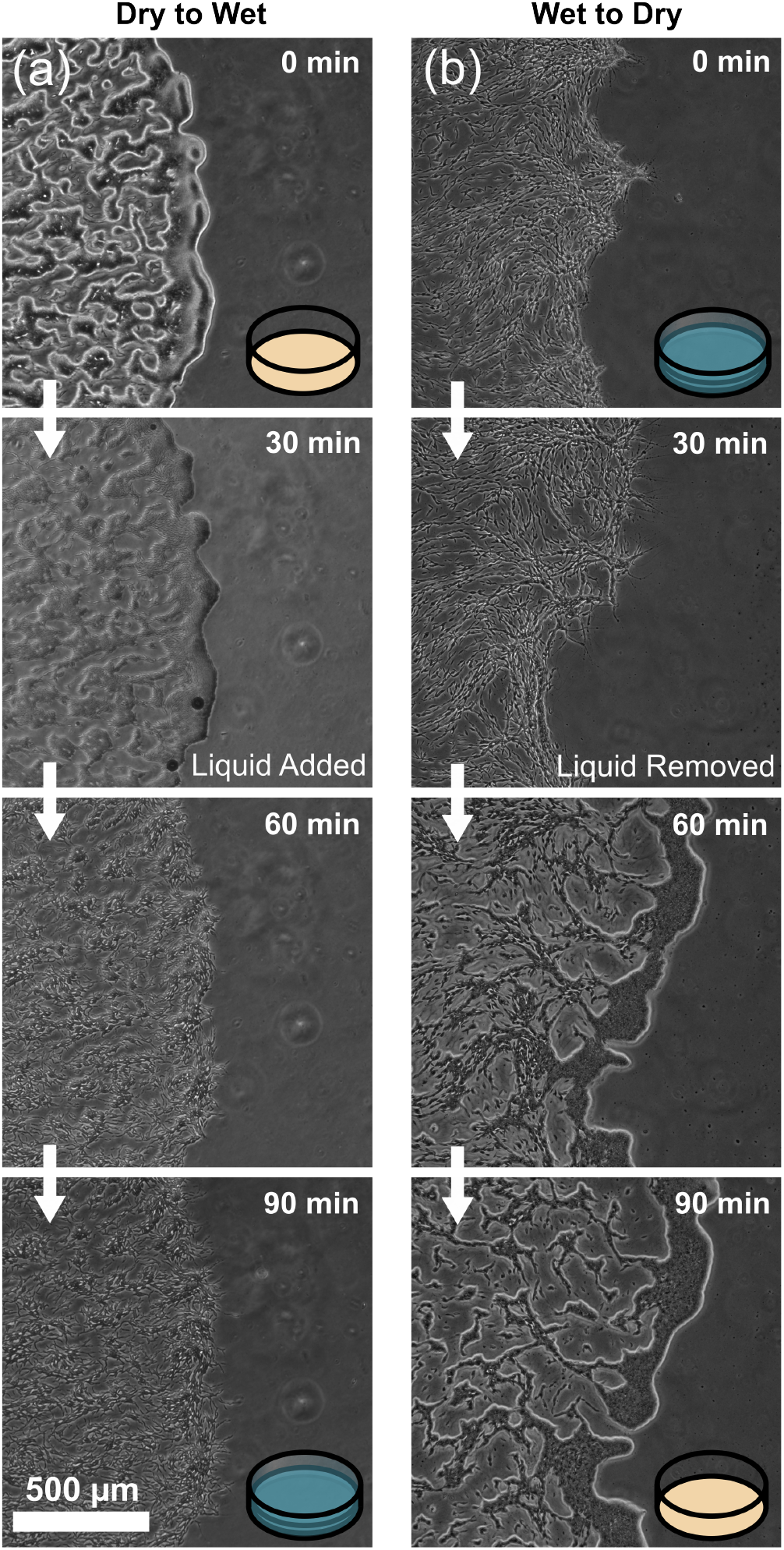
Morphological change with substrate immersion. (a) Addition of seawater to an initially air exposed, densely packed colony results in a restructuring to the network morphology. (b) Upon removal of the seawater overlay from an intially submerged colony, a restructuring from network to dense morphology is observed. Captured in phase contrast.

**FIG. 6.**
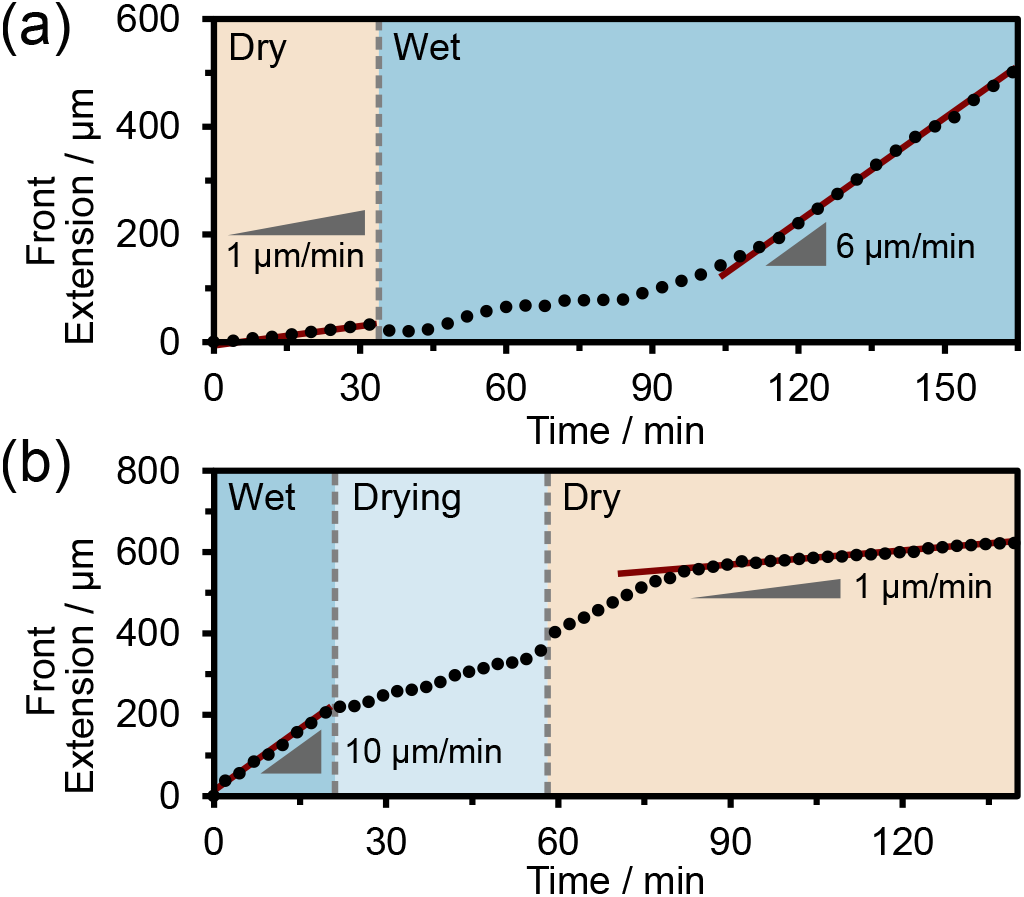
Change in Expansion Rate. (a) An initially air exposed colony sees an increase in expansion rate upon the addition of a seawater overlay. (b) For the reverse, a reduction in front propagation speed as the colony dries to an air interface. Rates calculated from increase in colony area between 30 s frame intervals.

To understand such morphological transition, we appeal to recent work on bacterial colonies [57]. Black et al. suggest that evaporation-induced shrinkage of the agar gel surface establishes an osmotic pressure difference *P*_0_ across the agar interface. So, a cell with a hydrophilic surface placed on the agar will be wetted to give a curved meniscus that generates a capillary pressure to balance *P*_0_. The capillary attraction generated by the overlapping menisci between nearby cells then gives rise to a densely-packed, nematic state [57]. For *Labyrinthula*, the capillary attraction between slime net trackways can then explain the observed compact colony morphology under dry conditions. No such capillary attraction exists between completely submerged trackways, which therefore adopt a sparse, network morphology characteristic of the genus.

### Development of network morphology

Next, we explore how a submerged slim-net morphology emerges in time by injecting cells from an axenic culture into an SSB immersion flowcell with plastic bottoms (recall that colony growth is substrate-independent, Fig. 4) and imaging subsequent development using phase-contrast microscopy. Our observations are summarised schematically in Fig 7(a).

**FIG. 7.**
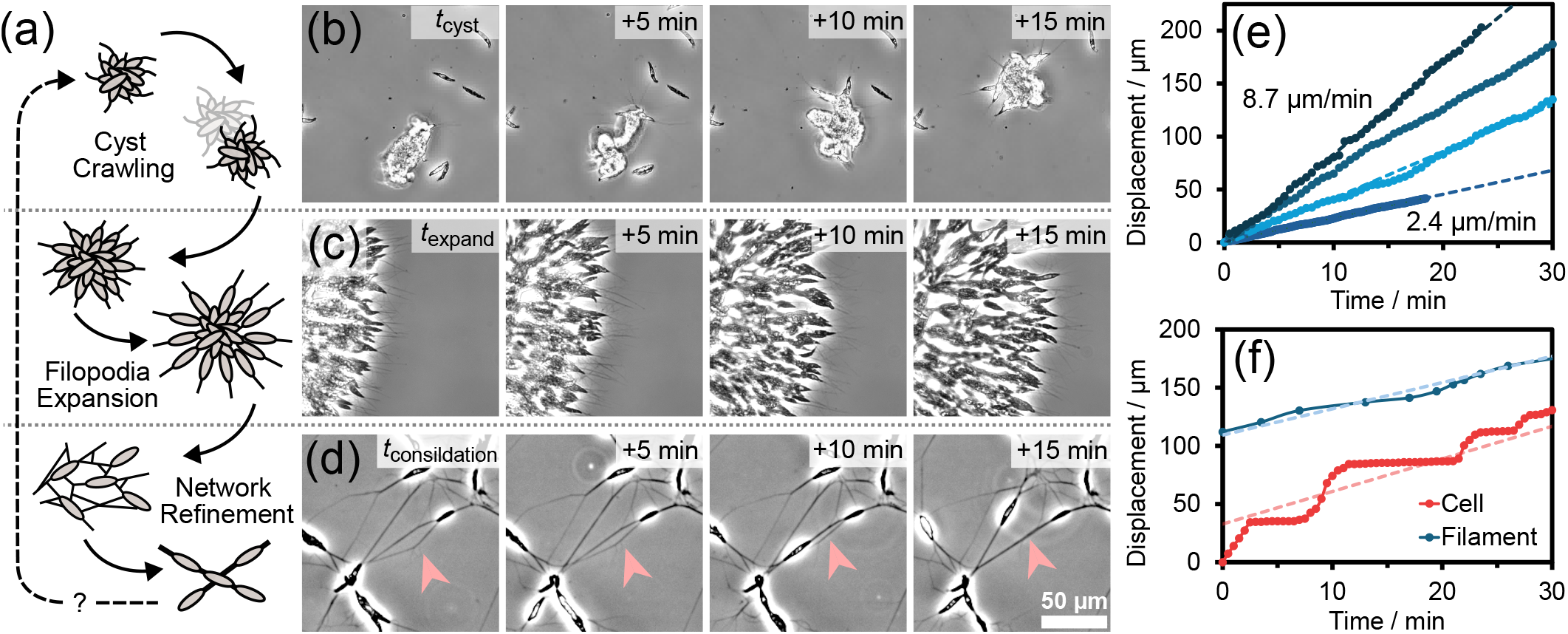
Filopodal-like filament extension. (a) Schematic representation of the typical features observed in the microscopic development of the network morphology, including tentative reversion to cyst structures from network consolidation. (b) Initial clumps of cells (cysts) are seen to migrate across the substrate surface within minutes of inoculation (*t*_cyst_ ∼30 min). (c) After some time *t*_expand_ ≳ 30 min, cysts appear fixed and radial filopodia extension is proceeded by outward cell motion through the growing filament ‘trackways’. (e) For a given filament, speed is seen to be constant during extension. Across colonies, extension speeds are seen to vary in the range 2 −10 µm min^−1^. (f) The displacement of a cell progressing through an extending filament shows an erratic profile with bursts of motion (shaded grey) interspersed with stationary periods resulting in an approximately constant separation to the filament tip. (d) Within the established network, filaments are seen to approach laterally and apparently fuse to form a single trackway.

#### Development from single cysts

The field of view immediately after inoculation shows a few medium-size clusters of many tens of cells and many smaller lenticular units, many of which appear to consist of a doublet of cells with phase-dark tips. Within minutes, the larger clusters engage in vigorous coordinated motion, Fig. 7(b) (Electronic Supplementary Video 2), somewhat reminiscent of the ‘migrating slug’ stage in the growth cycle of *Dictyostelium discoideum* [58]. During this phase of development, which has a typical timescale *t*_cyst_ ∼30 min, filopodia-like protrusions emerge from the periphery of these mobile cell clusters. At this stage, their main function does not appear to act as trackways for 2-dimensional cellular motility on the substrate. Instead, they seem to be ‘fishing’ for smaller clusters along and above the substrate surface (Electronic Supplementary Video 2, 3). After contacting a smaller cluster, a filopodium would then shorten to pull its ‘cargo’ into the main cluster. Such filament contractions also appear to be responsible for the bulk migration of cysts, effectively pulling them across the substrate. At these early times, trackways are evidently not permanently tethered to the substrate: they can be seen moving sideways, contracting, etc. Some of these space exploring filopodia undergo twisting motion, sometimes to form braids; a filopodium may also buckle while contracting (Electronic Supplementary Videos 4 and 5). Similar twisting and buckling has been studied in filopodia emerging from human cells as they explore the surrounding extracellular space [51], and may indicate that the actin filaments within the ectoplasmic membrane at this early stage adopt helical configurations [50].

#### Growth and restructuring of the network morphology

Eventually, a cluster will stop moving and enter a distinct radial expansion phase at *t*_expand_ ≳ 30 min, Fig. 7(c)). Thereafter, trackways appear to be locked in position, presumably by actin filament ‘guy ropes’ [42], and extend distally at speeds of 2 −10 µm min^−1^, Fig. 7(e). This is comparable to filopodia extension speeds observed in other organisms [59], but much lower than the ∼ 1 µm s^−1^ reported for *Labyrinthula* filaments by Preston et al. [42]; however, the latter appear consistent with speeds achieve in the ‘fishing’ motion described above, Supplementary Fig. S2.

During this stage of radial extension, the growing slimeways increasingly function as trackways proper, with cells moving bidirectionally along them. Such ‘traffic’ along membrane-bound trackways of actin shows striking similarities with an increasing body of observations concerning ‘tunnelling nanotubes’ (TNTs) found connecting mammalian cells that seem to fulfil a variety of functions, including the inter-cellular transport of viruses and organelles [60].

We observe cell motion along trackways at speeds in the range 1 −2 µm s^−1^, consistent with prior reports [42] and considerably faster than filopodia extension speeds. The latter discrepancy results in an erratic speed profile for cells at the expanding front of the colony, Fig 7(f). Here, cells progress radially outwards in bursts of motion, frequently overtaking temporarily motionless cells along a trackway. Interestingly, this erratic motion gives rise to an approximately constant separation between the tip of the extending filament and the nearest cell proceeding it, given by the offset between the red and blue dashed lines in Fig 7(f) (see also electronic supplementary material Figure S1). This is consistent with such ‘frontier cells’ playing a direct role in filament extension.

Over the course of hours, the radial expansion of colonies results in an extended network across the substrate of the flow-cell. Within this mature network, trackways reconstruct and consolidate from *t*_consolidation_ ≳ 2 h. For example, trackways may approach laterally and apparently fuse into a single new track, Fig 7(d). The functional significance of such network restructuring remains unclear. Tentatively, it may represent a spatial optimisation of the overall network structure, potentially enhancing cellular transport or nutrient uptake, similar to the restructuring observed in the slime mold *Physarum* [61]. In the longer term, such restructuring may be implicated in the reformation of cyst-like structures in the absence of nutrients [45].

### SUMMARY AND CONCLUSIONS

We have studied the growth and morphology of *Labyrinthula* colonies on a variety of solid substrates under different environmental conditions. The slime net colony form characteristic of the genus is only found when an overlying liquid layer is present, and transitions to a dense morphology when growing on a solid substrate exposed to air. One may speculate that such a transition occurs twice daily in intertidal seagrass meadows through cycles of submersion and drying. One intertidal species, *Z. noltii*, was largely unaffected [11] by the wasting disease epidemics in the 1930s that destroyed a large portion of North Atlantic seagrass [10]. Given that *Labyrinthula* is implicated in wasting disease, it is possible that the switch between network and dense forms of the organism’s colony growth may play a role in the distinctive response of *Z. noltii*.

The arrangement of ectoplasmic membranes in network colonies, Fig. 2, was clarified some time ago by high-resolution electron microscopy [38], but their arrangement in dense colonies has not been probed to date. It is perhaps possible that the inner ectoplasmic membranes of at least some neighbouring cells in a dense colony are fused, but the matter can only be resolved by future ultramicroscopy.

Our observation that *Labyrinthula* is able to develop into network colonies on submerged glass, plastic and agar substrates is consistent with the presence of the organism on a range of substrates in coastal habitats [45, 46, 48, 49], but with one caveat. In nature, the predominant feeding mode of the organism is likely through predation of other microbes [18], presumably facilitated by its slime net in a way that is still to be elucidated.

Although some structural [38, 41, 42] and biochemical [42] details are known, the biophysics of *Labyrinthula* motion along trackways remains unclear. While this mode of locomotion may well turn out to be unique, there are potential parallels to be drawn with the transport of organelles and other ‘cargo’ such as virions in tunnelling nanotubes (TNTs) [60], although different myosins are probably involved (myosin I in *Labyrinthula* [42] and myosin X in TNTs [60]). Our observations of the initial growth of the ectoplasmic network from cysts reveals twisting and buckling motion of filopodia as they explore space, recalling similar motion in filopodia emerging from human cells [51]. These parallels suggest that further studies of *Labyrinthula* biophysics may illuminate other areas of biology. Conversely, advances elsewhere should throw light on *Labyrinthula*. Thus, for example, quantifying how its filopodia twist and buckle and comparing these observations to theory [50] may give insights into the arrangement of actin filament within the outer ectoplasmic membrane and the filament-membrane interaction.

Finally, the function of the ectoplasmic net in *Labyrinthula* and how this unique feature relates to its potential for pathogenicity on seagrass remain unclear. To move towards such understanding, future work may explore, e.g., how the observations reported here alter in the presence of prey such as the diatom *Nitzschia* [19].

## Supporting information

Supplementary Movie 1 - Dense Phase Motion

Supplementary Movie 2 - Crawling Motion

Supplementary Movie 3 - Fishing Motion

Supplementary Movie 4 - Twisting Motion

Supplementary Movie 5 - Buckling Motion

Supplementary Movie 6 - Cell and Filament Dynamics Tracking

## Acknowledgements

We thank Dr Jane Polglase (Strameno Ltd.) for drawing our attention to the peculiar locomotion of *Labyrinthula*, and are grateful to Alastair Lyndon and Kirsty Fraser (Heriot Watt University) for teaching us how to collect the organism from seagrass. We also thank Dr Jochen Aarlt for helping us optimise our microscopy protocols, and Dr Lucas Le Nagard for many fruitful discussions.

