## Supplementary Information for "Environmental dependence of colony morphologies in *Labyrinthula* species"

#### 1. Erratic speed profile for cells behind expanding filaments

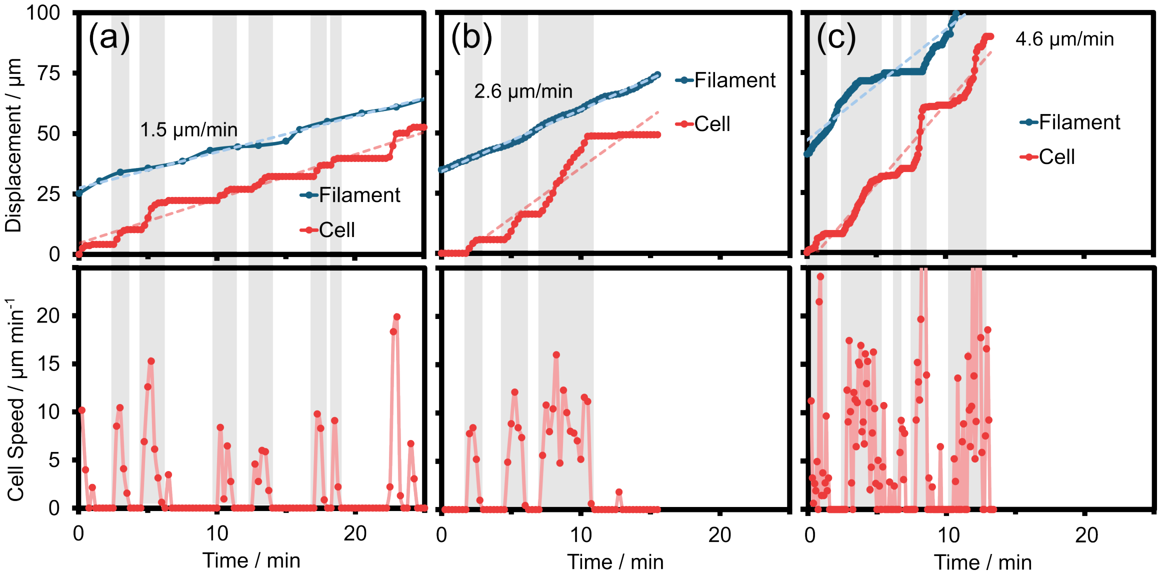


Figure S1: **Filament-Cell Extension Dynamics.** Across a variety of filament extension rates (a)-(c), the cells behind the extending filament exhibit an erratic speed profile whilst maintaining an approximately constant separation with the filament tip. Top panels show the cumulative displacement of filament tip and cell whilst bottom panels show corresponding instantaneous cell speeds between frames.

### 2. Filopodia fishing motion

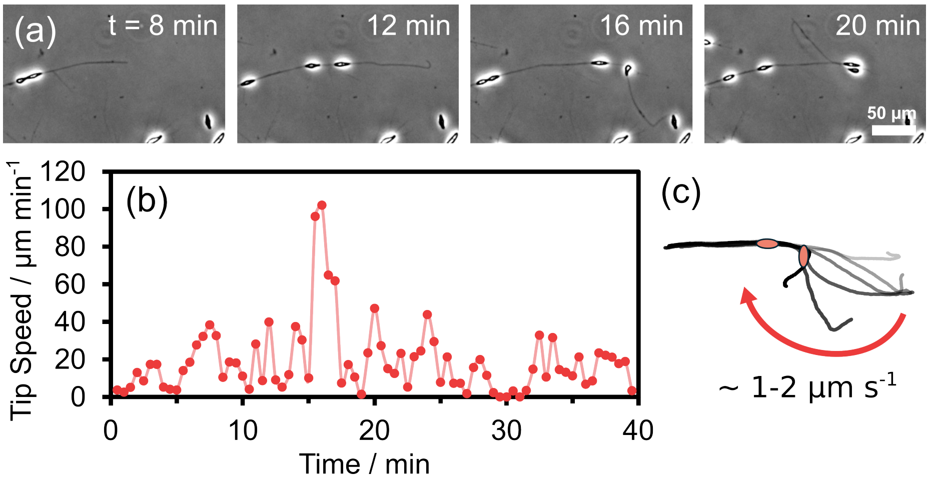


Figure S2: **Filament-Fishing Motion.** In early stages of colony development, filaments appear untethered from the substrate, describing sweeping ('fishing') motions with speeds in the $\sim 1\,\mu\mathrm{m}\,\mathrm{s}^{-1}$ range. (a) Snapshots from Supplementary Video 3. (b) Corresponding frame-to-frame speed. (c) Schematic of the motion.

### Supplementary Video 1 - Dense Phase Motion

Mature *Labyrinthula* colony (20 hours) grown on 1.2% agar Seawater Serum Agar (SSA) plate with no liquid overlay. Cell motion is observed in this dense phase. Timelapse captured in brightfield with a 10x air objective.

### Supplementary Video 2 - Crawling Motion

Crawling of cell clumps on the lower substrate of the flowcell in the minutes following inoculation. Ectoplasmic net filaments are seen to emanate from the cell clump and appear to facilitate the motion of the clump. Captured in phase contrast with a 10x air objective.

### Supplementary Video 3 - Fishing Motion

An extending ectoplasmic net filament with cell following is seen to detach from the substrate and performing a sweeping motion partially out of the plane of focus, resulting in the filament crossing back over itself. Capture in phase contrast with a 10x air objective.

### Supplementary Video 4 - Twisting Motion

High time resolution capture showing apparent twisting and braiding of ectoplasmic net filaments. Captured in phase contrast with a 10x air objective.

### Supplementary Video 5 - Buckling Motion

Apparent retraction of a filament appears to result in buckling, suggestive of a helical actin structure [1]. Captured in phase contrast with a 10x air objective.

### Supplementary Video 6 - Cell and Filament Dynamics Tracking

Showcase of the erratic motion of a cell following an extending filament. Extension tracking of the cell and filament tip is performed using TrackMate [2]. Displacement between tracked coordinates $\Delta(t_i, t_j) = \sqrt{(x_j - x_i)^2 + (y_j - y_i)^2}$ is evaluated to calculate cumulative displacement for plots. Captured in phase contrast with a 10x air objective.
